# The genomic landscape of recombination rate variation in *Chlamydomonas reinhardtii* reveals a pronounced effect of linked selection

**DOI:** 10.1101/340992

**Authors:** Ahmed R. Hasan, Rob W. Ness

**Affiliations:** Department of Cell and Systems Biology, University of Toronto, Toronto, ON M5S 3G5, Canada; Department of Biology, University of Toronto Mississauga, Mississauga, ON L5L 1C6, Canada

**Author notes:** Correspondence: Ahmed Hasan.

**Keywords:** recombination rate variation, linked selection, frequency of sex, GC-biased gene conversion, *Chlamydomonas*

## Abstract

Recombination confers a major evolutionary advantage by breaking up linkage disequilibrium (LD) between harmful and beneficial mutations and facilitating selection. Here, we use genome-wide patterns of LD to infer fine-scale recombination rate variation in the genome of the model green alga *Chlamydomonas reinhardtii* and estimate rates of LD decay across the entire genome. We observe recombination rate variation of up to two orders of magnitude, finding evidence of recombination hotspots playing a role in the genome. Recombination rate is highest just upstream of genic regions, suggesting the preferential targeting of recombination breakpoints in promoter regions. Furthermore, we observe a positive correlation between GC content and recombination rate, suggesting a role for GC-biased gene conversion or selection on base composition within the GC-rich genome of *C. reinhardtii*. We also find a positive relationship between nucleotide diversity and recombination, consistent with widespread influence of linked selection in the genome. Finally, we use estimates of the effective rate of recombination to calculate the rate of sex that occurs in natural populations of this important model microbe, estimating a sexual cycle roughly every 770 generations. We argue that the relatively infrequent rate of sex and large effective population size creates an population genetic environment that increases the influence of linked selection on the genome.

## Introduction

Recombination, the shuffling of existing genetic material, is both a fundamental evolutionary process and required to ensure proper disjunction of chromosomes during meiosis. Meiotic recombination has two possible outcomes: crossing over (CO) and non-crossing over (NCO), also known as gene conversion. There is clear evidence that recombination rate varies at multiple scales across nature, with variability observed within and between taxa as well as within the genome (Stapley et al., 2017). Recombination reduces interference between linked adaptive and harmful mutations and therefore is an important determinant of how well natural selection can act. The spatial heterogeneity of recombination within genomes means that local recombination rate is an important determinant of the efficacy of selection and other evolutionary processes in a given genomic region (Hill and Robertson, 1966, Felsenstein, 1974, McVean et al., 2004). Understanding the predictors of recombination rate variation has thus constituted a core objective in the study of how recombination affects genome evolution.

Across the species that have been examined to date, predictors of recombination rate variation remain inconsistent (reviewed in Choi and Henderson 2015, Stapley et al. 2017). In many mammalian genomes, which display a high degree of fine-scale variability in recombination rate, the presence of recombination hotspots is known to be driven by the double strand break-inducing histone H3 methyltransferase protein PR domain-containing 9 (PRDM9) (Baudat et al., 2010, Parvanov et al., 2010, Baudat et al., 2013). However, studies of recombination in non-mammalian model organisms have uncovered conservation of hotspot regions over long evolutionary time scales at or near functional genomic elements, such as near transcription start sites (Tsai et al., 2010, Singhal et al., 2015, Lam and Keeney, 2015). Hotspots in *Saccharomyces cerevisiae* have been found to preferentially occur in GC-rich promoter regions (Gerton et al., 2000, Lam and Keeney, 2015) and regions of open chromatin (Wu and Lichten, 1994, Berchowitz et al., 2009). Yet other predictors are less consistent in their effects on recombination frequency across systems. For example, local methylation is observed to suppress recombination in *Arabidopsis thaliana* (Yelina et al., 2015) yet in humans, germline methylation levels are positively correlated with recombination rate (Sigurdsson et al., 2009). Similarly, GC content is positively correlated with recombination rate in humans and yeast, but not in *A. thaliana* (Fullerton et al., 2001, Mancera et al., 2008, Wijnker et al., 2013). Finally, in contrast to the hotspot-punctuated recombination landscapes observed in the above species, the *Caenorhabditis elegans* genome is largely devoid of hotspots altogether, instead displaying relatively large genomic blocks of constant recombination rate (Rockman and Kruglyak, 2009, Andersen et al., 2012). Whilst the presence of PRDM9 does explain many of the patterns observed in mammalian genomes, a variety of other predictors have been observed thus far, often without consistent effects between systems.

High-resolution estimates of genome-wide recombination rate variation have thus far been constrained to a few model animals, fungi, and land plants, which represent a very small subset of eukaryote diversity. What determines recombination landscapes and whether PRDM9-like drivers of recombination rate exist in other taxa remains to be discovered. The recombination profiles of most protists remain unknown, and in some of those cases, whether they are even sexual in the first place (D'Souza and Michiels, 2010, Grimsley et al., 2010, Stapley et al., 2017). It is estimated that the rate of sex is unknown in nearly all (>99%) free-living protist species (Weisse, 2008). The rate of sex is especially relevant to overall estimates of recombination in the case of unicellular eukaryotes that switch between clonal and sexual reproduction. In a primarily asexual population, it follows that mutation will generate the vast majority of novel genetic material over evolutionary time, while the relative reduction of recombination will, all else being equal, result in a more pronounced effect of linked selection (Agrawal and Hartfield, 2016, Hartfield, 2016). The rate of sex can be estimated as the relative frequency of meioses to mitoses by combining direct estimates of the recombination (*r*) and mutation (*μ*) rate with population estimates of genetic diversity (4*N_e_μ*) and the effective recombination rate (4*N_e_r*) (Tsai et al., 2008). Although this estimation is possible, there are very few organisms in which diversity (*θ*), LD (*ρ*), recombination rate (*r*), and mutation rate (*μ*) are known.

Here, we examine recombination rate variation across the genome in the facultatively sexual model green alga *Chlamydomonas reinhardtii*, and estimate its rate of sex in nature. We use a population genomic method to infer recombination rtes, allowing for a more fine-scale assessment than would otherwise be possible with traditional linkage mapping. In this study, we use whole-genome sequencing of 19 *C. reinhardtii* field isolates from two nearby localities in Quebec, Canada, to address the following questions using LD-based approaches: 1) What is the landscape of recombination rate variation in the genome of *C. reinhardtii* ? 2) What genomic features predict recombination rate variation? 3) How does recombination rate correlate with diversity? 4) What is the rate of sex in natural populations of *C. reinhardtii* ? In doing so, we present a fine-scale estimate of genome-wide recombination rate variation in the organism, and find an enrichment of recombination hotspots immediately upstream and downstream of genes; a pattern consistent with observations made in other non-mammalian systems.

## Results

### The variable landscape of recombination in *C. reinhardtii*

From our population of 19 individuals (Table S1), we calculated fine-scale recombination landscapes across the genome of *C. reinhardtii* using the coalescent-based population recombination rate estimator LDhelmet 1.7 (Chan et al., 2012). LDhelmet estimates recombination rate between adjacent pairs of SNPs in units of *ρ*/bp, where *ρ* = 2*N_e_r*; this unit is henceforth referred to as *ρLD*. The genome-wide average recombination rate was *ρLD* = 0.0044/bp, while mean recombination rates for each chromosome varied over an order of magnitude from 0.0026 to 0.0115, inversely scaling with chromosome lengths (Fig. S2, *R*^2^ = 0.4803, p = 0.002). Genome-wide *ρLD* estimates were then summarized in non-overlapping 2 kb windows for fine-scale analysis, with windows showing variation in recombination rate of up to an order of magnitude; the distribution of recombination rates is shown in Fig. 1A. We then investigated for the presence of recombination hotspots, defining a hotspot as a region that was 1) >2 kb in length and 2) exhibited a >5-fold increase in *ρLD* as compared to the surrounding 80 kb of sequence on either side, following previous hotspot definitions (Singhal et al., 2015). Under these criteria, we found evidence of hotspots in all chromosomes, with 904 hotspot regions in total representing 2.65% of the genome, where the average *ρLD* within hotspots was more than 6 times the genome average (mean length = 3.19 kb, mean *ρLD* fold increase over local background = 10.09, mean distance between adjacent hotspots = 123 kb, mean *ρLD* at hotspots = 0.02799).

**Figure 1:**
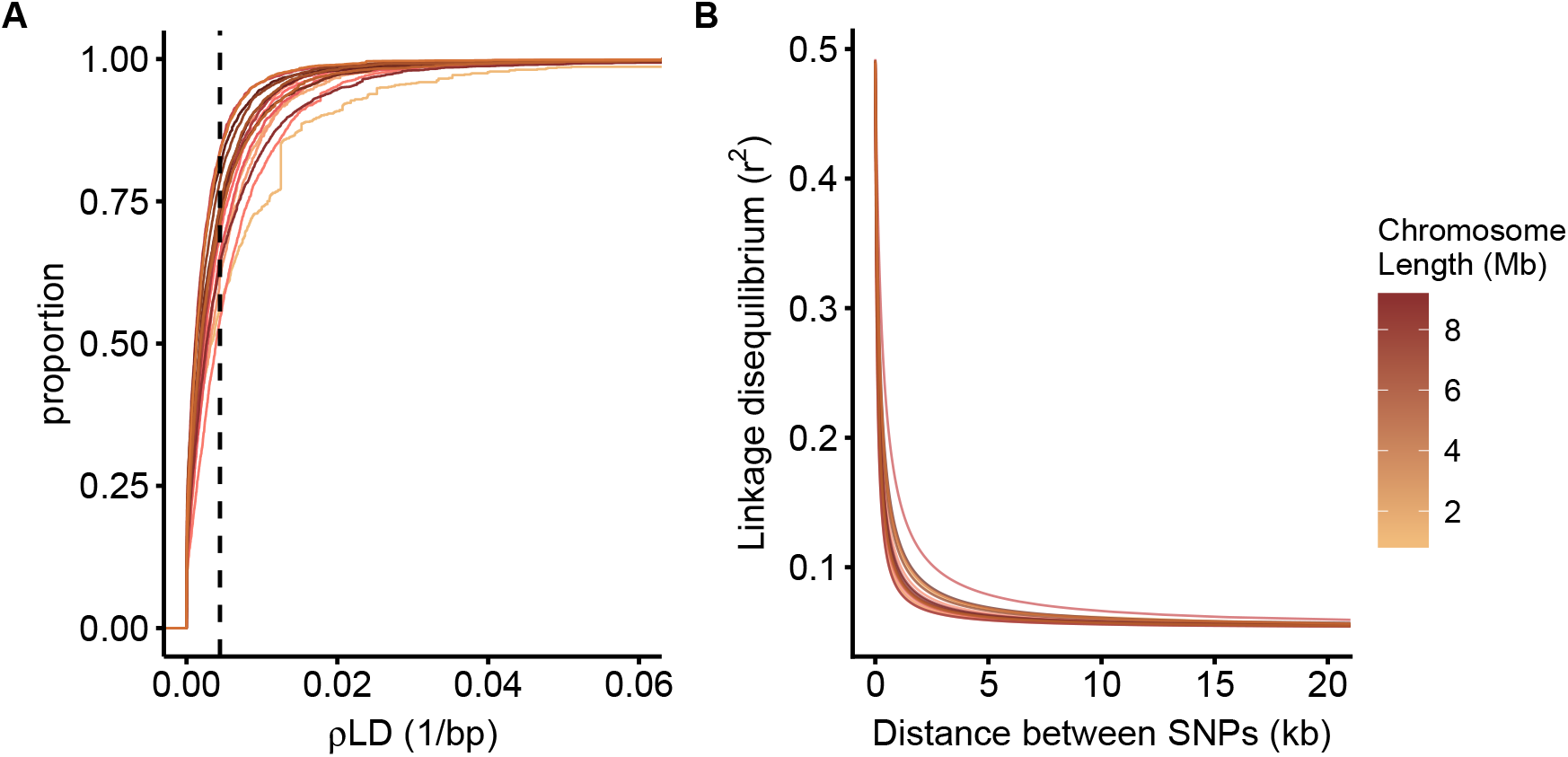
**A** Cumulative frequency distribution of *ρLD* for each chromosome of *C. reinhardtii*. Each curve represents one of the 17 chromosomes, shaded by chromosome length. *ρLD* values were summarised in 2 kb windows. The vertical dashed line indicates the genomewide mean *ρLD* value. **B** Decay of linkage disequilibrium (*r*^2^) across the 17 chromosomes of *C. reinhardtii*, modeled according to equation 3 from Hill and Weir (1988).

To examine the decay of LD with physical distance in *C. reinhardtii*, we calculated pairwise LD (*r*^2^) between SNPs on each chromosome using plink v1.90 (Chang et al., 2015). We then modeled the expected decay of LD with physical distance for each chromosome following equation 3 from Hill and Weir (1988) (Fig. 1B). Estimates of LD measured as *r*^2^ dropped to half of their starting value within a mean distance of 164 bp (range 93 - 394 bp). Moreover, the decay of *r*^2^ leveled off within a mean distance of 9500 bp, where ‘leveling off’ was defined as the point at which the instantaneous rate of change in *r*^2^ with physical distance approximates 0 to six significant digits (mean *r*^2^ at level off point = 0.0585 *±* 0.0017). In concordance with the relationship observed between LDhelmet *ρLD* values and chromosome length, we also observed that LD approaches baseline levels faster in shorter chromosomes, although this correlation is marginally nonsignificant (Spearman’s *ρ* = 0.4559, p = 0.067).

### Recombination rate is highest immediately surrounding genes

To investigate the correlates of recombination across the genome, we examined how *ρLD* varied with different functional annotations in the *C. reinhardtii* reference genome (Merchant et al., 2007) (Fig. 2). We found that recombination rate was 11.9% higher in intergenic regions (mean *ρLD* = 0.004877) than genic regions (mean *ρLD* = 0.04358, Mann-Whitney U test, p *<* 2.2 *×* 10^*−*16^). Within intergenic regions, proximity to genes is a strong predictor of elevated recombination rate, with sites within 2 kb of genes displaying 24.2% higher mean *ρLD* than the genome background (mean *ρLD* of sites within 2 kb of genes = 0.005510). There is a striking and statistically significant 63% increase in recombination within the 2 kb upstream of genes compared to the adjacent 5’ UTR (Mann-Whitney U test, p = 3.16 *×* 10^*−*16^; *ρLD* upstream = 0.005707, *ρLD* 5’ UTR = 0.003498). Contrary to our expectations, protein-coding sequence had significantly higher rates of recombination than either the 5’ UTR or introns (Mann-Whitney U test, p = 5.69 *×* 10^*−*5^; *ρLD* CDS = 0.004838, *ρLD* introns = 0.004211). Finally, we examined hotspot enrichment across annotations, finding highest enrichment in the 2 kb of sequence flanking genes (Fisher’s exact test, p < 2.2 *×* 10^*−*16^). 13.9% of all hotspots occur in the 8.7% of the genome corresponding to these annotations.

**Figure 2:**
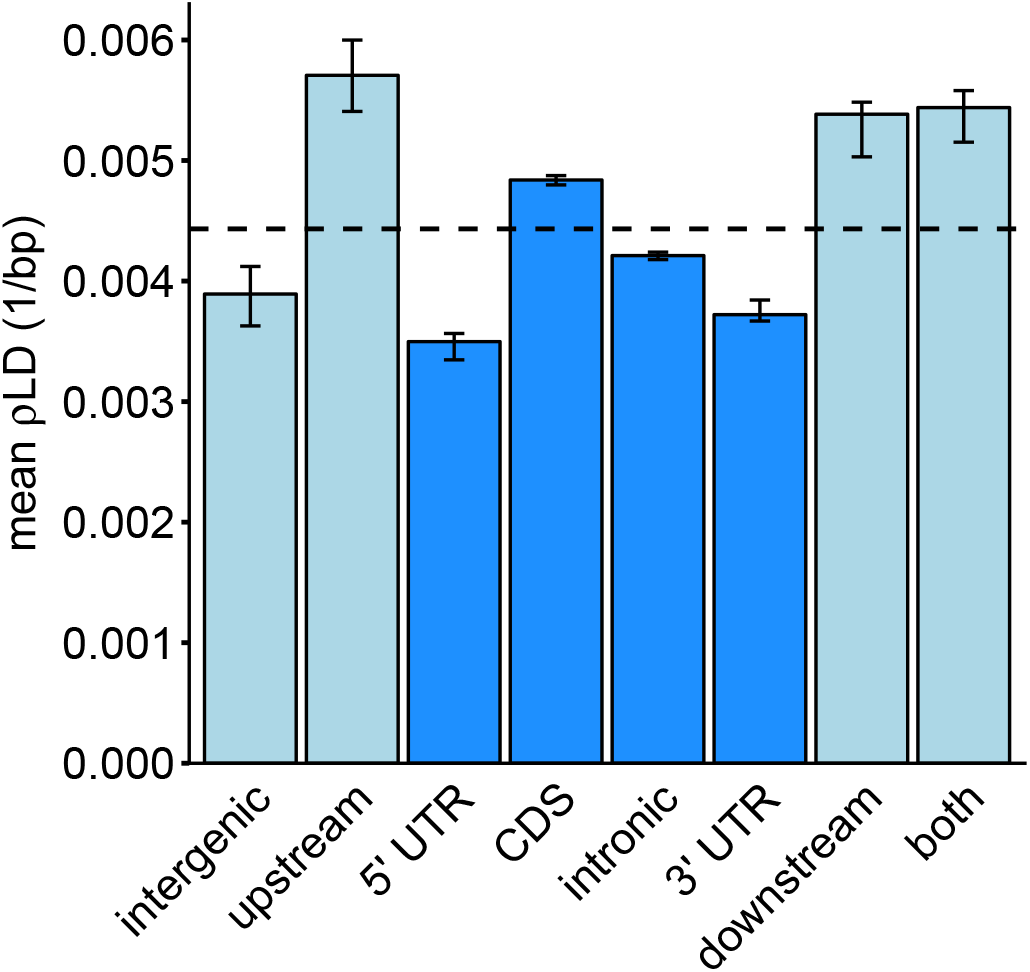
*ρLD* in different annotation categories across the *C. reinhardtii* genome, with intergenic annotations in light blue and genic annotations in dark blue. Error bars represent bootstrapped 95% confidence intervals (n = 1000). The annotations ‘upstream’ and ‘downstream’ represent intergenic sites that were within 2 kb of a genic region, while ‘both’ indicates sites that are downstream of one gene and upstream of another. ‘intergenic’ sites are all other intergenic sites more than 2 kb from the nearest gene. The dashed horizontal line represents the mean genome-wide *ρLD* value.

The association between recombination rate and genome annotation has been hypothesized to be driven by differential patterns of DNA methylation across functional annotations. Sequence methylation is subdivided into three types, depending on the sequence context surrounding a methylated cytosine: CG, CHG, and CHH (where H = A, T, or C). Given that the enrichment of recombination hotspots flanking genes in *C. reinhardtii* suggests a role for open chromatin in determining recombination localization (see Discussion), the open chromatin model would thus predict recombination suppression in heterochromatic regions (Choi and Henderson, 2015). To examine the relationship between DNA methylation patterning and re-combination rate in *C. reinhardtii*, we obtained bisulfite sequencing data from three clones of one of the Quebec *C. reinhardtii* strains included in the present analysis (CC-2937) (Kronholm et al., 2017), and summarised methylation levels across all three contexts and *ρLD* values in 200 bp windows. Here, we opted for a much smaller window size than in previous analyses because we expected that the highly localized nature of DNA methylation (Suzuki and Bird, 2008) would mean that effects on recombination would be at very fine scales. We found that the correlation between *ρLD* and methylation, although significant, is only very weakly negative (Spearman’s *ρ* = −0.044, p *<* 2.2 *×* 10^*−*16^), indicating that DNA methylation is not a strong driver of recombination in *C. reinhardtii*.

### Recombi;nation rate is positively correlated with nucleotide diversity

If background selection and selective sweeps are common, we expect that linked selection will drive a reduction in neutral diversity in regions of low recombination. To examine whether linked selection was occurring in the *C. reinhardtii* genome, we first examined the relationship between our *ρLD* estimates and neutral nucleotide diversity (*θ_π_* = 4*N_e_μ*) in non-overlapping 10 kb windows using a partial Spearman’s correlation. Here, we controlled for functional density, defined as the number of coding sites per window, since functional density may correlate with either or both of recombination rate and nucleotide diversity and confound the relationship between the two (Payseur and Nachman, 2002, Flowers et al., 2012). We found *θ_π_* at selectively unconstrained (intronic, intergenic, and four-fold degenerate) sites was positively associated with *ρLD* (*R*^2^ = 0.3385, p *<* 2.2 *×* 10^*−*16^), in alignment with expectations from linked selection (Fig. 3A). Concordant with this observation, we found nucleotide diversity to be nearly 50% higher in hotspot regions (*θ_π_* = 0.02676, 95% CI = 0.0265-0.0273, p *<* 2.2 *×* 10^*−*16^) than in coldspots, or regions exhibiting 5-fold *ρLD* reductions relative to the surrounding 80 kb of sequence (*θ_π_* = 0.01793, 95% CI = 0.0179-0.0181, p *<* 2.2 *×* 10^*−*16^).

**Figure 3:**
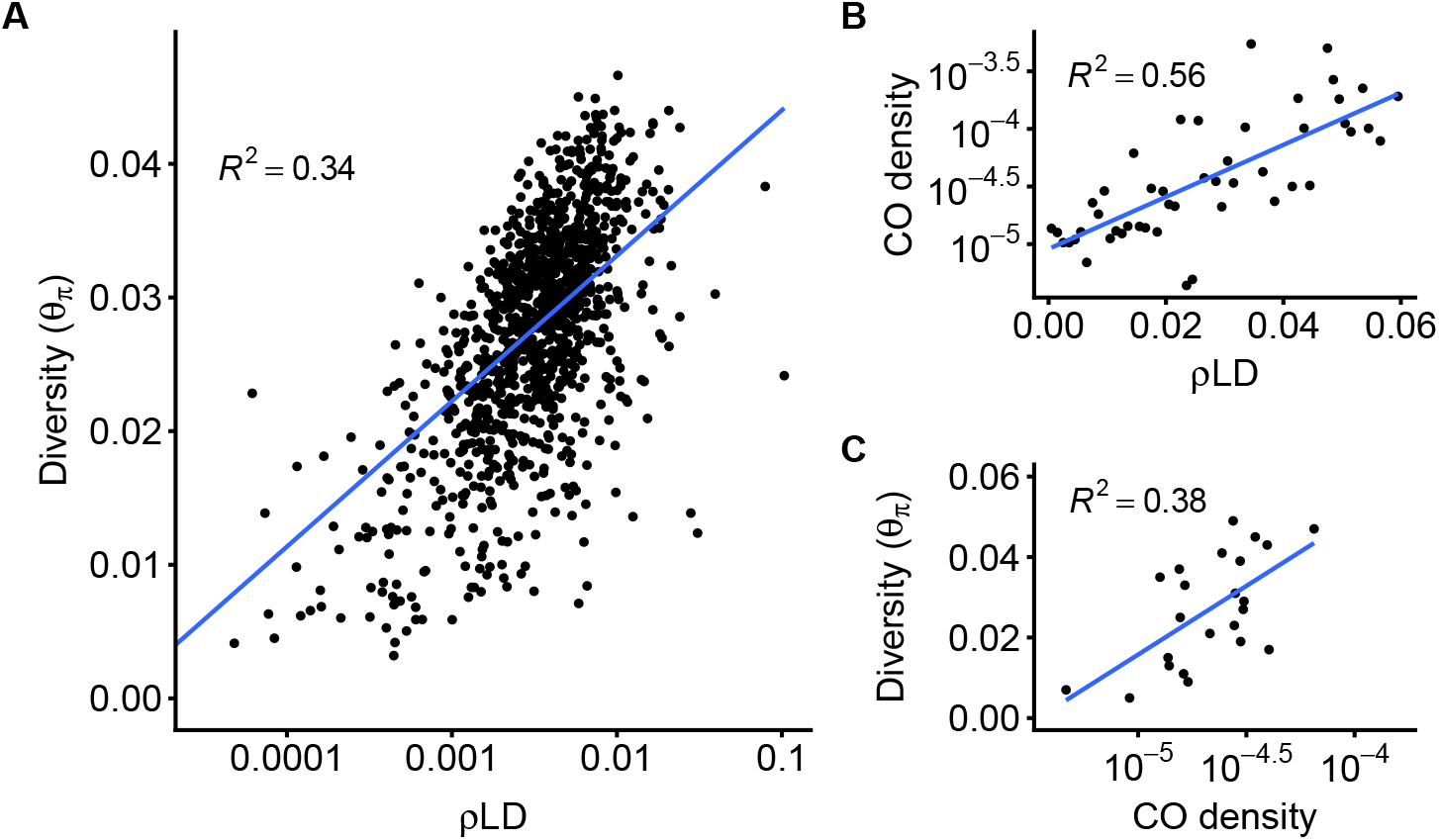
**A** Neutral diversity (*θ_π_*) scales with LD recombination rate in 100 kb windows. Nucleotide diversity was calculated at intronic, intergenic, and 4-fold degenerate sites. For clarity, we have constrained the plot to windows where *ρLD* was greater than 5 × 10^*−*4^. **B** LD recombination rate is correlated with crossover density, as obtained from the dataset of Liu et al. (2017). **C** Crossover density from the Liu et al. (2017) dataset correlates with neutral diversity.

Despite the strong correlation observed above, it stands that both diversity and *ρLD* are confounded by *N_e_*, which will vary across the genome due to selection, linkage, and variation in local coalescence time. To test whether the patterns observed using *ρLD* reflects physical recombination rate *r* and not just variance in *N_e_*, we obtained the crossover data of Liu et al. (2017), who sequenced 27 F1 tetrads obtained from a cross between *C. reinhardtii* strains CC2935 *×* CC2936 (both of which are included in the present analysis) and directly inferred the locations of recombination events from sequence data. We binned *ρLD* estimates from genomic intervals, and then for each bin, calculated CO density as the number of COs that occured in genomic regions with that *ρLD* value over the total number genomic sites with that range of *ρLD* values. This measure of CO density was then used as a proxy for physical recombination rate, which is unaffected by *N_e_*. Here, we obtained a positive and significant correlation between *ρLD* and log-scaled CO density, weighting for the number of sites in each bin (Fig. 3B, *R*^2^ = 0.559, p = 9.84 *×* 10^*−*10^), indicating that variation in *ρLD* is indeed reflective of variation in *r* and not just *N_e_*.

Next, we extended the same binning method to directly investigate whether COs tend to occur more frequently in regions of the genome with higher diversity. Here, we calculated *θ_π_* in windows across the genome and then binned sites according to their diversity. (see Methods). We calculated CO density values for the sites in the genome corresponding to each bin. We found CO density and *θ_π_* were also significantly correlated (Fig. 3C, *R*^2^ = 0.383, p = 0.0016), further suggesting the action of linked selection in the genome.

### Recombination rate is positively correlated with GC content

Recombination rate has also been found to correlate with GC content in a variety of systems (Mugal et al., 2015). This relationship is often attributed to GC-biased gene conversion (gBGC), in which heteroduplex mismatches between GC and AT bases formed during recombination are preferentially repaired in favour of GC nucleotides (Galtier and Duret, 2001). From a population genetic perspective, this effect is indistinguishable from positive selection in favour of GC alleles, and can thus substantially impact genome evolution (Duret and Galtier, 2009, Ratnakumar et al., 2010). *C. reinhardtii* is known to have a highly GC-rich genome (64%) despite a strong A/T mutational bias, raising the possibility that gBGC may have played a role in the evolution of its genome (Ness et al., 2015). Here, we find that GC content does display a positive correlation with recombination rate in non-overlapping 10 kb windows (Spearman’s *ρ* = 0.3028, p *<* 2.2 *×* 10^*−*16^).

### Estimating the frequency of sex in *C. reinhardtii*

Due to the fact that mutations arise every cell division (each meiosis and mitosis) yet only a fraction of cell divisions are sexual (*f*) and allow recombination to occur. We can therefore use estimates of neutral diversity (*θ* = 2*N_e_μ*) and population recombination rate (*ρ* = 2*N_e_r*) combined with lab estimates of *μ* and *r* to roughly estimate the relative frequency of mitosis to meiosis, or the frequency of clonal relative to sexual reproduction (Ruderfer et al., 2006, Tsai et al., 2008). If we are to take *r* to represent *r* multiplied by the frequency of sex *f* relative to clonal reproduction (such that *ρ* = 2*N_e_rf*), we can use existing estimates of the other parameters to solve for the ratio 1*/f*.

Thus, our genome-wide recombination rate estimate of 4.43 *×* 10^*−*3^ can be used in tandem with previous estimates of the *C. reinhardtii* recombination rate (*r* = 12 cM/Mb) (Liu et al., 2017), the mutation rate (*μ* = 9.63 *×* 10^*−*10^) (Ness et al., 2015), and neutral diversity (*θ* = 2.75 *×* 10^*−*2^) (Ness et al., 2016) to estimate the frequency of sex in natural populations of *C. reinhardtii*. With this approach, we obtain *f* = 0.001292, corresponding to one sexual generation for every 1*/f* = ~ 770 asexual generations.

## Discussion

In this study, we have generated a fine-scale estimate of the recombination rate landscape in *C. reinhardtii* using patterns of linkage disequilibrium in whole-genome resequencing data, revealing a punctuated recombination landscape with frequent hotspots. We also find an enrichment of recombination hotspots within 2 kb of genes that leads to an overall increase in recombination rate surrounding genes, in concordance with observations in a number of other PRDM9-lacking organisms. The variation in recombination rate across the genome is correlated with nucleotide diversity, suggesting that the influence of linked selection is widespread in the genome and that recombination is a major driver of genetic variation. Lastly, we have used our estimate of the population recombination rate to estimate the frequency of sex as being once every ~ 770 generations in *C. reinhardtii*, allowing for further insights into the ecology and evolution in of this model organism in nature.

Assuming an *N_e_* of 1.4 *×* 10^7^ (Ness et al., 2016), the genome-wide *ρLD* estimate of 4.43 *×* 10^*−*3^ can be converted to an estimate of *r* = *ρ/*2*N_e_* = 0.016 cM/Mb, following the method described in Auton and McVean (2012). Although this estimate of genome-wide *r* on its own is approximately two orders of magnitude below most plants (Henderson, 2012), including the parameter *f* (= 0.001292) such that *r* = *ρ/*2*N_e_f* yields a genome-wide recombination rate estimate of *r* = 12.25 cM/Mb, which is closer to the estimate of *r* = 9.15 cM/Mb from the genetic map of *C. reinhardtii* (Kathir et al., 2003, Merchant et al., 2007, Henderson, 2012). This suggests that in *C. reinhardtii*, infrequent sex reduces effective recombination, bearing consequences for the effects of linked selection in the genome (see below).

Between chromosomes, we observe variation in mean recombination rates across an order of magnitude, and also find that recombination rate inversely correlates with chromosome length (Fig. S2), a relationship consistent with prior studies in a variety of organisms (Kaback et al., 1992). Given that each chromosome requires at least one crossover event to ensure proper meiotic disjunction (Page and Hawley, 2003), it follows that shorter chromosomes exhibit higher per-base crossover rates, resulting in more pronounced signatures of LD breakdown. Furthermore, *C. reinhardtii* exhibits moderate rates of LD decay across all 17 chromosomes. Our estimates of the distance (*≤*10 kb) at which LD (*r*^2^) decays to baseline LD levels are shorter than those obtained in a previous study of *C. reinhardtii* that reported a decay to baseline within ~ 20 kb (Flowers et al., 2015). The difference between our estimates may be caused by genetic structure in *C. reinhardtii*; where we sequenced isolates all from nearby localities, Flowers et al. utilized a mix of lab strains alongside isolates from a variety of populations across Quebec and Eastern United States. This disparity in our respective estimates supports the idea that there are barriers to recombination across the geographic range of *C. reinhardtii* in North America, with the resulting population structure increasing LD among variants.

At the 2 kb scale, we find widespread recombination hotspots across the genome, similar to observations in mammals, angiosperms, and yeast (Stapley et al., 2017). On the other hand, this recombination profile is unlike that of both *C. elegans*, which has a comparatively homogenous fine-scale recombination landscape (Rockman and Kruglyak, 2009, Kaur and Rockman, 2014) as well as *D. melanogaster*, which displays some degree of fine-scale heterogeneity but little evidence for highly localized elevations in recombination rate (Chan et al., 2012, Manzano-Winkler et al., 2013).

We see elevated recombination and an enrichment of hotspots within regions immediately flanking genes in *C. reinhardtii*, consistent with a trend conserved across the lineages of nearly all PRDM9-lacking organisms studied thus far. Specifically, recombination hotspots upstream of genes have been observed in fungi (Berchowitz et al., 2009, Tsai et al., 2010, Lam and Keeney, 2015), finches (Singhal et al., 2015), as well as angiosperms, such as wheat (Saintenac et al., 2009), maize (Li et al., 2015) and *A. thaliana* (Wijnker et al., 2013, Choi et al., 2013). In addition, the same pattern is observed in the mammalian Canidae family, where PRDM9 was lost relatively recently (Axelsson et al., 2012, Auton et al., 2013). In these PRDM9-lacking organisms, chromatin structure has been invoked as an explanation of recombination hotspot conservation upstream of genes (Wu and Lichten, 1994, Lichten, 2008, Berchowitz et al., 2009). In eukaryotes, DNA is wrapped in nucleosomes, with nucleosome occupancy depleted in regions where the DNA needs to be accessible, such as for the purposes of transcription. Promoter regions upstream of genes thus tend to display greater nucleosome depletion, which may in turn allow for recombination machinery to more easily induce double strand breaks in these regions (Pan et al., 2011, Lam and Keeney, 2015). Our observations of elevated recombination rate immediately flanking genes suggest a similar mechanism acting in the *C. reinhardtii* genome, and show that this trend is even more wide-spread, extending to green algae.

Within genes, however, we observe more surprising patterns in recombination rate variation. We might expect that recombination rates will evolve to be lower in functional regions, so as to suppress adverse mutagenic effects that may result from crossing over. Furthermore, recombination events in functional regions that result in deleterious mutations may be selected against and therefore reduce evidence of recombination. The 5’ UTR in *C. reinhardtii* displayed the lowest recombination rate of any annotation, despite recombination rate elevations observed in the UTRs of other species (Mancera et al., 2008, Kawakami et al., 2017). One explanation could be that recombination-associated deleterious mutations in the functional components of the UTR region might have been eliminated by selection, increasing observed LD. However, were this the case, we would expect a similar trend in protein-coding exons, where we instead observe the highest rates of within-gene recombination. Moreover, recombination rate in intronic sequence is intermediate to the UTR and exonic sequence, whereas under a mutagenic hypothesis we would instead expect within-gene recombination rates to be higher in introns than in exons. Altogether, the idea that selection has reduced effective recombination in functional sites does not appear to hold from the current data. Thus, either recombination is not mutagenic in *C. reinhardtii*, or we must invoke another mechanism, such as chromatin conformation or some yet to be discovered driver of recombination, to explain the observed patterns. Unfortunately, relatively little is known about chromatin conformation in *C. reinhardtii*, and we see only a very weak association with methylation, leaving identification of the drivers of recombination rate variation in and around genes an open question.

We find that *ρLD* positively correlates with nucleotide diversity across the genome of *C. reinhardtii*, indicating the action of linked selection (Fig. 3). This correlation is also observed using the CO dataset of Liu et al. (2017) as a proxy for physical recombination rate (Fig. 3C), which unlike *ρLD* and *θ_π_* is not scaled by *N_e_*. Theory predicts that the correlation of recombination and diversity arises as a consequence of background selection and/or selective sweeps reducing diversity in regions of otherwise low recombination (Charlesworth et al., 1993, Cutter and Payseur, 2013, Campos et al., 2017). We also find diversity to be higher in hotspot regions than coldspot regions, which is to be expected if linked selection is less pronounced in regions of elevated recombination.

Our result suggests that linked selection is a strong determinant of standing genetic variation in *C. reinhardtii*. Given that *C. reinhardtii* is likely to have a very high effective population size (Ness et al., 2012) it is expected that many mutations will be effectively selected (i.e. *N_e_s >* 1) (Kimura, 1962). However, while the effective population size is very large, the relatively infrequent rate of sex (see below) means that the effective recombination rate is not particularly high relative to obligately sexual species. The interaction of a large *N_e_* facilitating efficient selection alongside reduced recombination due to a facultatively sexual life cycle means that the influence of linked selection may be pronounced in the genome, and modulated less by recombination rate *per se* than would be the case in obligate outcrossers. This principle may be more widespread in unicellular eukaryotes, which live in large populations that are only periodically sexual.

In addition, we also find a weakly positive correlation between GC content and local recombination rate. Our result is consistent with a trend seen in other organisms such as yeast (Gerton et al., 2000), mouse (Jensen-Seaman et al., 2004), and humans (Fullerton et al., 2001) (although *A. thaliana* is a notable exception, displaying an inverse correlation - see Wijnker et al. 2013). The correlation we observe is in line with the presence of GC-biased gene conversion (gBGC) (Galtier and Duret, 2001) acting to drive up GC content in the *C. reinhardtii* genome. The potential action of gBGC is especially notable considering that despite a strong A/T mutational bias, *C. reinhardtii* has a GC-rich genome (64.1%) (Merchant et al., 2007, Ness et al., 2015). While our result lends support to a possible role of gBGC, a recent study revealed only weak gBGC from the genome sequences of 27 *C. reinhardtii* tetrads, in concert with a low overall rate of gene conversion, thus indicating a very minor role for biased gene conversion in the evolution of its genome (Liu et al., 2017). It is worth noting that using LD-based estimates of population recombination rate, we obtain a stronger correlation between GC content and recombination rate (Spearman’s *ρ* = 0.2434) than Liu et al. (Spearman’s *ρ* = 0.0646). A stronger GC-recombination correlation when considering historical recombination events suggests that the effects of weak forces governing base composition are more apparent over longer evolutionary timescales. However, there remain other possible explanations for the correlation between recombination and GC content past gBGC: first, simply that more recombination events initiating in GC rich regions, and second, more efficient selection for GC content in regions with higher recombination (Kliman and Hey, 1993, Campos et al., 2012).

Finally, by integrating lab- and population-based measures of recombination and mutation, we have estimated the rate of sex in *C. reinhardtii* to be one meiosis every ~770 asexual generations. The frequency is higher than estimates in *S. cerevisiae* (~50000 generations) (Ruderfer et al., 2006) as well as *S. paradoxus* (~ 1000-3000 generations) (Tsai et al., 2008). However, this method for estimating the frequency of sex is subject to numerous assumptions, chief amongst them neutrality and demographic equilibrium. We assume random mating, no gene flow, and no population subdivision; violations of these assumptions may otherwise result in more prominent LD and thus downwardly bias estimates of the frequency of sex. Despite these caveats, our estimate of sex occurring every ~ 770 generations may point towards a seasonal ecology in *C. reinhardtii*. While the precise rate of mitosis in nature is unknown, log-phase cultures subjected to a light dark cycle typically exhibit 2-3 doublings every 24 hours (Bernstein, 1964, Jones, 1970, Harris et al., 2009). An average of 2.5 doublings per day corresponds to 770 generations every 308 days. Considering the fact that sex is induced when conditions worsen and zygotes are resistant to freezing and other environmental stressors (Morris et al., 1979, Harris et al., 2009), it is plausible that populations of *C. reinhardtii* in Quebec overwinter as zygotes, undergoing a sexual cycle approximately once per year.

Taken together, our results suggest that populations of *C. reinhardtii* maintain relatively large effective sizes even at small geographic scales. These large populations are subject to efficient selection that interacts with infrequent bouts of sexual reproduction to drive strong effects of linked selection in the genome. The genome also displays significant heterogeneity in recombination rate, with recombination highest in the regions flanking genes; however, *C. reinhardtii* has relatively high recombination in coding exons, which suggests yet to be described drivers of recombination in this species.

## Materials and Methods

### Strains, sequencing, and alignment

19 (9 mt+, 10 mt-) natural strains of *Chlamy-domonas reinhardtii*, sampled from Quebec, Canada, were obtained from the Chlamy-domonas Resource Center. For strains CC2935, 2936, 2937, and 2938, we obtained publicly available sequencing data (Flowers et al., 2015). These 19 strains are all sampled from two nearby localities in Quebec, and show no evidence of population structure (R.J. Craig, personal communication) (summarized in Table S1). Sequencing and alignment were performed as described previously (Ness et al., 2016). Briefly: we conducted whole-genome resequencing on the Illumina GAII platform at the Beijing Genomics Institute, and aligned 100bp PE reads with BWA 0.7.4-r385 (Li and Durbin, 2009). Since the *C. reinhardtii* reference genome is derived from an mt+ individual and does not include organelles, we appended the chloroplast genome, the mitochondrial genome, and the mt-locus to allow mapping of reads derived from these regions. The GATK v3.3 tools HaplotypeCaller and GenotypeGVCFs were then used to call SNPs and short indels, and stored in Variant Call Format (VCF) files (non default settings: `ploidy=1`, `includeNonVariantSites=true`, `heterozygosity=0.02`, `indel_heterozygosity=0.002`).

### Recombination landscapes and hotspots

To obtain chromosome-wide maps of recombination rate variation in the *C. reinhardtii* genome, we used LDhelmet 1.7 (Chan et al., 2012), which calculates fine-scale estimates of population recombination in intervals between adjacent SNPs. The coalescent-based approach of LDhelmet also allows for inferences of ancestral recombination rate variation. LDhelmet reports the population recombination rate *ρ* = 2*N_e_r* that reflects the size of the recombining population (*N_e_*) and the physical recombination rate (*r*, recombination events · bp^-1^ · generation^-1^).

The block penalty parameter in LDhelmet determines the level of smoothing of *ρLD* estimates along the chromosomes. To ascertain the block penalty that reflects real variation in *ρLD*, we computed average *ρLD* in non-overlapping 500 bp windows across the longest chromosome (12) of *C. reinhardtii* using block penalties of 10, 50, and 100. We then performed pairwise correlations between windowed *ρLD* values across different block penalties and found that the estimates were virtually identical, which indicates that at the scale of hundreds of bases, our results were not sensitive to block penalty (Fig. S3). Here, we report data using the default LDhelmet setting of 50. We used a population scaled mutation rate of 0.01 (Ness et al., 2015), and for each chromosome ran LDhelmet for 1,000,000 iterations following 100,000 iterations of burn-in. Default parameters were otherwise retained.

To detect recombination hotspots in LDhelmet’s *ρLD* estimates, we modified a Python script from Singhal et al. (2015) that summarises *ρLD* values in non-overlapping windows. Following Singhal et al., we initially defined hotspots as regions that 1) were at least 2 kb in length and 2) exhibited a mean 5-fold increase in *ρLD* as compared to the surrounding 80 kb of sequence, following previous approaches (International HapMap Consortium, 2005, Singhal et al., 2015).

### LD decay across chromosomes

Pairwise calculations of *r*^2^ within each of *C. reinhardtii*’s 17 chromosomes were conducted using plink 1.90 (Chang et al., 2015). For all pairs of SNPs, plink calculates LD statistics with a maximum likelihood approach described in (Gaunt et al., 2007). By default, plink filters out pairs of SNPs with an *r*^2^ value below 0.2; we disabled this filtering. We calculated *r*^2^ for all pairs of SNPs within 100 kb of one another, and modeled the expected decay of LD with distance for each chromosome with non-linear least squares regression in R (R Core Team, 2017) using equation 3 from Hill and Weir (1988).

### Genomic correlates of recombination rate

We classified the reference genome of *C. reinhardtii* with the following annotations: genic sites were subclassified as protein coding sequence (CDS), exons, introns, and/or UTRs, while intergenic regions were classified as being within 2 kb upstream of a gene (‘upstream’), 2 kb downstream of a gene (‘downstream’), flanked by genes within 2 kb on either side (‘both’), or more than 2 kb from the nearest gene (‘intergenic’). For each of the above features, average *ρLD* was calculated from every corresponding site in the genome. Recombination rates for each annotation were bootstrapped for 1000 replicates in order to obtain 95% confidence intervals.

We then examined the relationship between DNA methylation and recombination rate using publicly available bisulfite sequencing data from three clones of *C. reinhardtii* strain CC-2937 (Kronholm et al., 2017). For read mapping, we used BWA-meth (Pedersen et al., 2014) and called methylated cytosines in CG, CHG, and CHH contexts using biscuit-0.2.2 (Zhou, 2017). Following Kronholm et al., we set minimum base quality to 20 and minimum mapping quality to 60. We then used the output VCF file to summarise methylation beta values in 200 bp windows, and examined the correlation between *ρLD* and methylation at the 200 bp scale. We also examined the relationship between methylation and recombination within each of the three sequence contexts, but observed the same pattern as in the overall correlation.

For the correlation of GC content with recombination rate, we used a custom Python script (https://github.com/aays/2018-ld-paper/blob/master/antr_diversity_gen.py) to compute GC content in a given window, and then summarized GC content with *ρLD* values in non-overlapping 10 kb windows.

### Recombination and nucleotide diversity

Nucleotide diversity (*θ_π_*) was calculated at ‘silent’ sites (intronic, 4-fold degenerate and intergenic) in 10 kb windows for downstream correlation with *ρLD*. We performed a partial Spearman’s correlation between *ρLD* and diversity while controlling for functional density, defined as the proportion of sites in a window overlapping a protein-coding exon, using the R package ppcor. To examine nucleotide diversity in and out of hotspots, we calculated *θ_π_* in these regions across the genome in non-overlapping 2 kb windows, and split the resulting dataset using our hotspot classification from earlier. Windowed *θ_π_* estimates were bootstrapped for 1000 replicates in order to obtain 95% confidence intervals.

To test whether variation in *N_e_r* was reflective of *r* and not just *N_e_*, we obtained the crossover data of Liu et al. (2017), who sequenced 27 tetrad offspring (= 108 individuals). We binned our *ρLD* estimates for each genomic interval into 50 bins between *ρLD* values of 0 and 0.06, and for each bin, counted the number of Liu dataset COs found in regions of the genome corresponding to that range of *ρLD* values. This count of COs was converted to a density measure by dividing the number of COs in each bin by the number of sites falling within a the *ρLD* values defining a given bin. Next, to examine the relationship between diversity and CO density, we first calculated *θ_π_* across the genome in 10 kb windows, and then binned these values across 50 bins ranging from 0 to 0.1. Windows with less than 500 silent sites were discarded to reduce noise in diversity estimates and bins with less than 100000 total genomic sites were discarded because with so few COs in the dataset these bins tended to have zero COs. Then, we assigned COs to bins based on local diversity in the region bounded by each CO and calculated COs per site as above.

All statistical tests in this work were implemented using R 3.4.1 (R Core Team, 2017). For all figures in this work, we utilized the R package ggplot2 (Wickham, 2009).

## Acknowledgements

Short read data have been uploaded to the European Nucleotide Archive under study accession ERP109393. This work was supported by a Natural Sciences and Engineering Research Council (NSERC) Discovery grant (RGPIN/06331-2016) and Canadian Foundation for Innovation John R. Evans Leaders fund (35591) to RWN. We thank S.I. Wright and A.M. Moses for helpful discussions and suggestions and Brian Novogradac for facilitating computational support and HPCNODE1. Scripts used in this work can be found at https://github.com/aays/2018-ld-paper. We declare we have no competing interests.

## Supplementary Information

**Figure S1:**
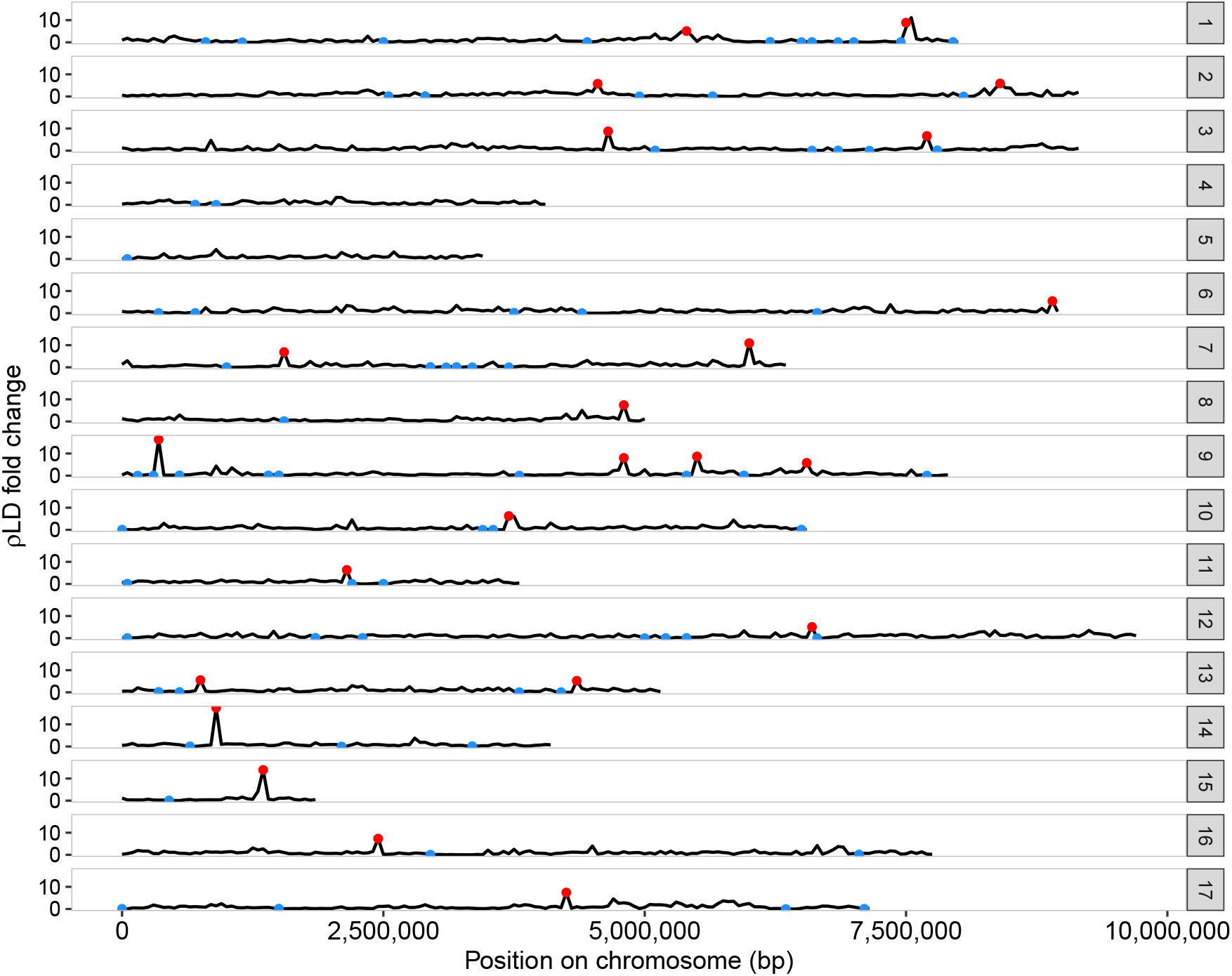
The landscape of recombination across the 17 chromosomes of *C. reinhardtii*, with *ρLD* fold change calculated against chromosomal background and summarised in non-overlapping 50 kb windows. Red dots represent regions displaying 5-fold elevations in recombination rate compared to chromosome background (‘hotspots’) whilst regions with 5-fold lower recombination than background (‘coldspots’) are marked in blue.

**Figure S2:**
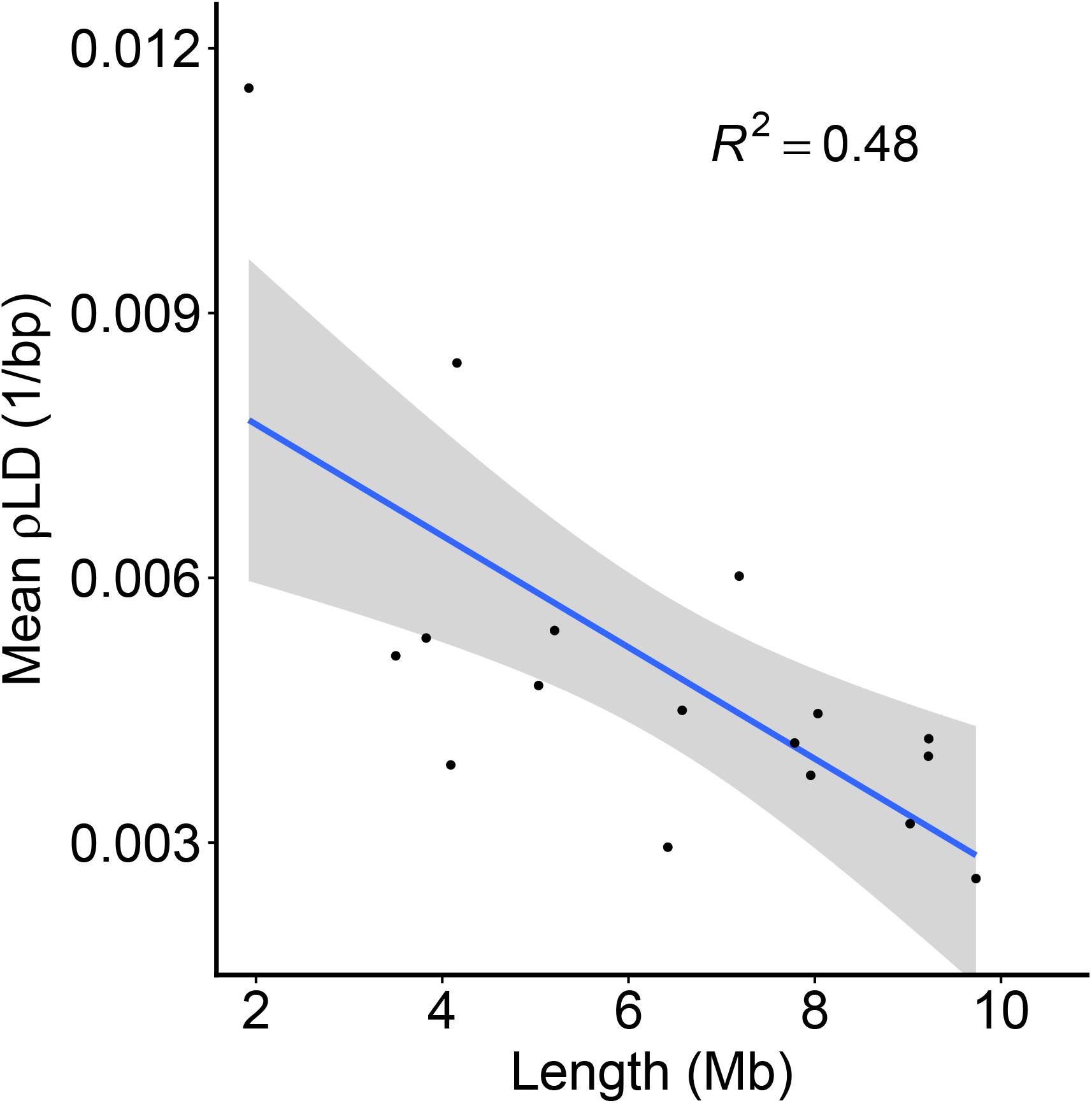
Chromosome length inversely scales with mean recombination rate (*R*^2^ = 0.4803). Each point represents one of the 17 *C. reinhardtii* chromosomes.

**Figure S3:**
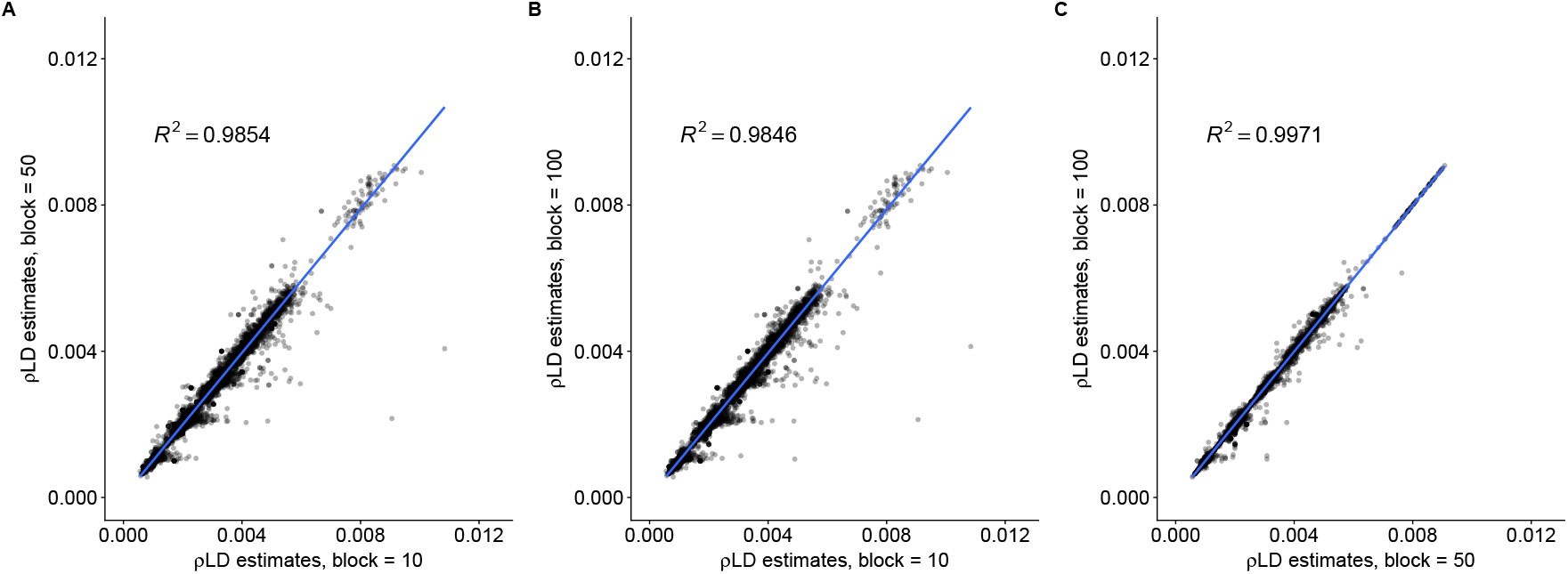
Variation in LDhelmet block penalty has minimal effects on recombination rate estimates. *ρLD* estimates from independent LDhelmet runs on chromosome 12 of *C. reinhardtii* over three different block penalties (10, 50, and 100) were summarised in non-overlapping 500 bp windows. Each point represents *ρLD* in one 500 bp window. **A** Comparison of *ρLD* estimates from LDhelmet runs at block = 10 and block = 50 (*R*^2^ = 0.9854) **B** Comparison of *ρLD* estimates from LDhelmet runs at block = 10 and block = 100 (*R*^2^ = 0.9846) **C** Comparison of *ρLD* estimates from LDhelmet runs at block = 50 and block = 100 (*R*^2^ = 0.9971).

**Table S1:** Field strains of *C. reinhardtii* used in this study. All strains were obtained from the *Chlamydomonas* Resource Center (chlamycollection.org). Mating types of mt-strains CC-3059 and CC-3062 are mislabelled as mt+ on the Resource Center website, and are instead mt-individuals (R.J. Craig, personal communication)

| Strain | Collection Location/Year | Mating type |
| --- | --- | --- |
| CC-2936 | Farnham, Quebec/1993 | + |
| CC-2937 | Farnham, Quebec/1993 | + |
| CC-3060 | Farnham, Quebec/1995 | + |
| CC-3064 | Farnham, Quebec/1995 | + |
| CC-3065 | Farnham, Quebec/1995 | + |
| CC-3068 | Farnham, Quebec/1995 | + |
| CC-3071 | Farnham, Quebec/1995 | + |
| CC-3076 | Montreal, Quebec/1995 | + |
| CC-3086 | Montreal, Quebec/1995 | + |
| CC-2935 | Farnham, Quebec/1993 | - |
| CC-2938 | Farnham, Quebec/1993 | - |
| CC-3059 | Farnham, Quebec/1995 | - |
| CC-3061 | Farnham, Quebec/1995 | - |
| CC-3062 | Farnham, Quebec/1995 | - |
| CC-3063 | Farnham, Quebec/1995 | - |
| CC-3073 | Farnham, Quebec/1995 | - |
| CC-3075 | Montreal, Quebec/1995 | - |
| CC-3079 | Montreal, Quebec/1995 | - |
| CC-3084 | Montreal, Quebec/1995 | - |

